# Oncogenic GNAQ/11-induced remodeling of the IP3/Calcium signaling pathway protects Uveal Melanoma against Calcium-driven cell death

**DOI:** 10.1101/2024.11.25.625282

**Authors:** Garcia Céline, Roussel Louis, Massaad Sarah, Brard Laura, La Rovere Rita, Tartare-Deckert Sophie, Bertolotto Corine, Bultynck Geert, Leverrier-Penna Sabrina, Penna Aubin

## Abstract

Despite being considered a rare tumor, uveal melanoma (UVM) is the most common adult intraocular malignancy. With a poor prognosis and limited treatment options, up to 50% of patients develop metastases, primarily in the liver. A range of mutations and chromosomal aberrations with significant prognostic value has been associated with UVM pathogenesis. The most frequently mutated genes are GNAQ and GNA11, which encode the α subunits of Gq proteins and are described as driver mutations that activate multiple signaling cascades involved in cell growth and proliferation. Directly downstream of Gαq/11 activation, PLCβ engagement leads to sustained production of DAG and IP3. While the DAG/PKC/RasGRP3/MAPK signaling branch has been identified as an essential component of UVM unregulated proliferation, the role of IP3-mediated signals has been largely overlooked.

Here, we demonstrate that, whilst maintaining Ca²⁺ homeostasis, UVM cells have developed a decoupling mechanism between IP3 and ER Ca²⁺ release by altering IP3 receptor (IP3R) expression. This correlation was observed in human UVM tumors, where IP3Rs were found to be downregulated. Critically, when IP3R3 expression was restored, UVM cells exhibited an increased tendency to undergo spontaneous cell death and became more sensitive to pro-apoptotic modulators of IP3R-mediated Ca²⁺ signaling, such as staurosporine and the Bcl2-IP3R disrupter peptide BIRD2.

Finally, inhibition of the Gαq/11 signaling pathway revealed that IP3R expression is negatively regulated by GNAQ/11 oncogenic activation. Hence, we demonstrated that by remodeling IP3R expression, GNAQ/11 oncogenes protect UVM cells against IP3-triggered Ca²⁺ overload and cell death. Therefore, the GNAQ/11 pathway not only drives proliferation through DAG activity but also provides a protective mechanism to evade IP3/Ca²⁺-mediated cell death. These dual functions could potentially be exploited in novel combinatorial therapeutic strategies to effectively block UVM cell proliferation while simultaneously sensitizing them to cell death.

## INTRODUCTION

Uveal Melanoma (UVM) is the most frequently occurring intraocular malignancy in adults. Despite enucleation or radiotherapy treatments, 45-50% of UVM patients ultimately succumb to metastasis (1). Although skin cutaneous melanoma (SKCM) and UVM both derive from melanocytes, they are intrinsically distinct diseases showing remarkable differences (2). The *BRAF* mutations typically found in 60% of SKCM cases (and targeted by approved targeted therapies) are rare in UVM. Instead, nearly 90% of UVM are characterized by somatic gain-of-function mutations in guanosine nucleotide-binding protein Q (*GNAQ*) or its paralogue guanosine nucleotide-binding protein alpha-11 (*GNA11*) gene, which encode two highly homologous α subunits of Gq heterotrimeric proteins involved in mediating G protein-coupled receptors (GPCR) signaling (1). These driver mutations, almost exclusively occurring at position Q209, abolish their intrinsic GTPase activity. This causes a prolonged activation of the Gα_q_ and Gα_11_ subunits, which accumulate in a GTP-bound state (3, 4). Constitutively active Gα_q/11_ directly binds and activates phospholipase C β (PLCβ) which converts phosphatidylinositol 4,5-bisphosphate (PIP2) into diacylglycerol (DAG) and inositol triphosphate (IP3) (5).

Other recurrent oncogenic mutations in UVM are targeting either the cysteinyl leukotriene receptor 2 (CysLTR_2_), a GPCR (6, 7), or the downstream effector of the *GNAQ/11* signaling, PLCβ4. Although these driver mutations are mutually exclusive, they all lead to constitutive activation of the Gα_q/11_-coupled GPCR signaling, with a shared outcome of sustained DAG/IP3 production (7).

Downstream DAG production, the Ras activator Ras Guanyl Releasing Protein 3 (RasGRP3) is activated by phosphorylation by DAG-activated Protein Kinase C (PKC). RasGRP3 serves as a critical signaling node, linking oncogenic *GNAQ/11* to the RAS/RAF/MEK/ERK signaling pathway (8, 9). This highlights the DAG/PKC/RasGRP3/MAPK signaling branch as an essential component driving the uncontrolled proliferation of UVM cells.

Surprisingly, and despite the frequent use of IP3 and its derivatives as a readout for oncogenic activation in UVM, the impact of the sustained production of IP3 has been overlooked in that context. In non-excitable cells, one of the most common routes of calcium (Ca^2+^) signal generation is the production of IP3 *via* the engagement of PLC-coupled transmembrane receptors. IP3 binds to and opens its receptors (IP3R), acting as ER transmembrane Ca^2+^ channels mediating the release of ER-sequestrated Ca^2+^. The resulting depletion in the ER-Ca^2+^ content is sensed by the stromal interaction molecules (STIM) that relocate at ER-plasma membrane junctions to bind and open Orai Ca^2+^ channels (10, 11). The resulting influx of Ca^2+^, called store-operated Ca^2+^ entry (SOCE), replenishes the ER-Ca^2+^ content but also signals several physiological and pathological functions in cells. In malignant cells, Ca²⁺ homeostasis and signaling are often remodeled, contributing to tumor progression.

Since *GNAQ/11* oncogenic mutations lead to the constitutive generation of IP3 through sustained PLCβ activation (7), a constant release of ER-stored Ca^2^, combined with constitutive SOCE stimulation, is anticipated. Yet, the resting Ca^2+^ concentration and fluxes must be tightly regulated at all times, as any massive elevation of intracellular Ca^2+^ would inevitably cause cell injury or death by perturbing normal cell signaling including critical ER and mitochondrial functions (12–14). Despite isolated evidence of possible dysregulated Ca^2+^ signaling in UVM (15, 16), any specific GNAQ/11-induced alterations in Ca^2+^ homeostasis or signaling, or the functional consequences of such remodeling on UVM transformation and progression, remain unreported.

Hence, knowing that the constitutive activation of Gα_q/11_ leads to a continuous production of IP3, we hypothesized that *GNAQ/11-mutant (GNAQ/11 ^mut^*) UVM cells have developed protective mechanisms to circumvent a massive Ca^2+^ entry while preserving Ca^2+^ homeostasis to sustain proliferation. In this study, we demonstrate that oncogenic *GNAQ/11^mut^* UVM implement profound adjustments to their “Ca^2+^ signaling toolkit”, notably *via* a fine-tuned regulation of IP3R, to elude a lethal IP3-driven Ca^2+^ overload while maintaining functional Ca^2+^ signals for survival.

## MATERIALS AND METHODS

### Cell culture and reagents

92.1 (17) and MDA-MB-231 cells were purchased from ECACC and ATCC, respectively. MP41 and MM66 cell lines (18) were kindly provided by Dr S. Roman Roman (Curie Institute, Paris), M229 and M238 (19) by Dr A. Ribas (University of California), SU-DHL4 by Dr L. Bresson-Bepoldin (University of Bordeaux) and OCM-1 (20) by Dr C. Bertolotto (University of Côte d’Azur). Table 1 summarizes cell lines characteristics.

**Table 1.**
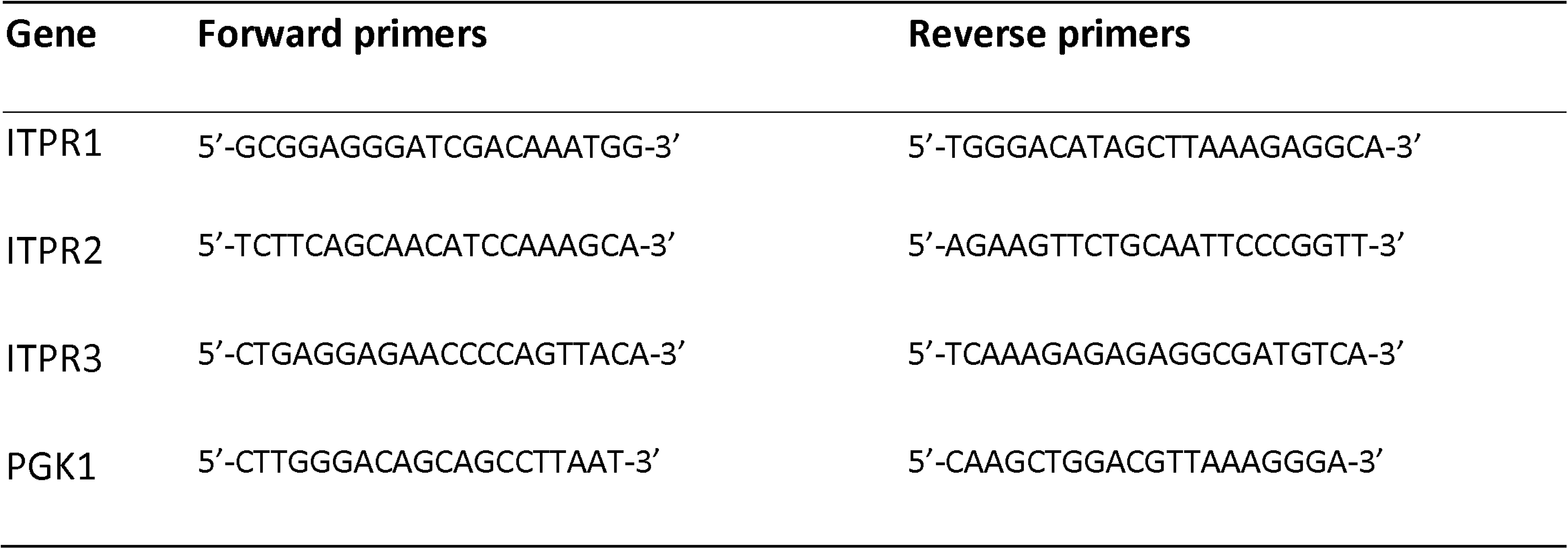
Cell lines characteristics. Summary of the origin, the driver and secondary genomic mutations, as well as the chromosomal alterations characterizing the different cell lines used in this study. PDX: Patient-derived xenograft. LOH: Loss of heterozygosity.

92.1, OCM-1, MP41, SU-DHL4 cells were cultured in RPMI 1640 (61870010, Fisher Scientific, Illkirch, France) supplemented with 10% Fetal Calf Serum (FCS) (CVFSVF0001, Eurobio, Les Ulis, France), or with 20% FCS for MM66 cells. M229, M238 and MDA-MDA-231 were cultured in DMEM (31966021, Fisher Scientific) with 10% FCS.

Thapsigargin (Tg) (586005), ionomycin (I0634) and staurosporine (569397) were purchased from Sigma Aldrich (St-Quentin Fallavier, France), AEB071 (S2791) from Selleckchem (Planegg, Germany), YM-254890 (AG-CN2-0509) from Coger (Paris, France) and were resuspended in DMSO. TAT-CTL (RKKRRQRRRGGSIELDDPRPR) and TAT-BIRD2 (RKKRRQRRRGGNVYTEIKCNSLLPLAAIVRV) peptides were synthesized by LifeTein (Hillsborough, NJ, USA) and resuspended in DMF.

### Cell transfection

For siRNA, cells plated at low density were transfected with 10 nM siRNA (control siRNA (siCTL) sequence is 5’-GCCGACCAAUUCACGGCCG-3’ and siRNA directed against *GNA11* (siGNA11) is 5’-AAAGGGUACUCGAUGAUGC-3’) using Lipofectamine™ RNAiMAX (13778150, Fisher Scientific) and harvested 48 h post-transfection.

For DNA plasmids, 5.10^6^ cells were electroporated with 10 μg of plasmids using the Gemini X2 generator (BTX Instrument Division, Harvard Apparatus, Holliston, MA, USA) and harvested 48 and 72h later. The mammalian eGFP-C1-IP3R3 (GFP-IP3R3) plasmid was kindly provided by Dr M. Pagano from the Howard Hughes Medical Institute.

### mRNA preparation and quantitative RT-PCR

Total RNAs were isolated using the NucleoSpin RNA plus kit (740984, Macherey Nagel, Hoerdt, France) following manufacturer’s instructions. mRNA concentrations were measured using the NanoDrop analyser ND100 (Labtech, Domont, France). cDNAs were obtained using the PrimeScript RT-PCR kit (RR037A, Takara, Saint-Germain-en-Laye, France) following manufacturer’s instructions. qPCR were performed by mixing cDNA (5 ng), primers (3 μM) and Takyon Low Rox SYBR Master Mix dTTP blue (UF-LSMT-B0701, Eurogentec, Angers, France) using the 7500 Fast Real-Time PCR System thermocycler (Applied Biosystems™, Foster City, CA, USA). Levels of expression were analyzed using the 2^-ΔCt^method when not compared to a control condition or using the 2^-ΔΔCt^. All values were normalized to the housekeeping gene, phosphoglycerate kinase 1 (PGK1) (ΔCt) and relativized to the control condition when needed (ΔΔCt).

List of primers sequences.

### Western blot assays

Cell pellets were lysed on ice in RIPA lysis buffer (89900, Fisher Scientific) supplemented with protease inhibitors (P8340, Sigma Aldrich) and phosphatase inhibitors (P5726, Sigma Aldrich). After sonication and 10 000 g centrifugation at 4°C, supernatants were dosed with the BCA kit (23227, Fisher Scientific) according to the manufacturer’s instructions. 30 µg of proteins were separated on 3-8% tris-acetate gels (10184482, Fisher Scientific), and transferred onto nitrocellulose membranes. Primary antibodies list: IP3R1 (75-035, Antibodies Inc, Davis, CA, USA); IP3R2 (sc-398434, Santa Cruz Biotechnology, Heidelberg, Germany); IP3R3 (610312, BD Biosciences, Le Pont de Claix, France); p-ERK (4370) and ERK (9102, Cell Signalling, Ozyme); HSC70 (sc-7298, Santa Cruz); a-tubulin (T6199, Sigma). Proteins were visualized using ECL RevelBlot Intense (OZYB002-1000, Ozyme, Saint-Cyr-L’Ecole, France) and quantified with the ImageJ software (NIH, Bethesda, MD, USA).

### Intracellular Calcium measurement in cell populations

Cells seeded in black 96-well clear bottom plates (736-0230, VWR, Rosny-sous-Bois, France) were loaded with 4 μM Fura-2AM (F1201, Fisher Scientific) for 45 min in OptiMEM (11564506, Fisher Scientific) or with the no quench Calcium 6 probe (R8190, Molecular Devices, Workingham, UK) for 2 h. Intracellular Ca^2+^ variations were measured using a multimode plate reader (Flexstation3, Molecular Devices) and analyzed with the SoftMax Pro software (Molecular Devices).

#### Mn^2+^ quenching

After Fura-2AM loading, cells were washed in a Ca^2+^-free solution (135 mM NaCl, 5.4 mM KCl, 5.6 mM glucose, 0.8 mM MgCl_2_, 10 mM HEPES, pH 7.4). Fluorescence emission was measured at 510 nm after an excitation at 360 nm. After a 35 s baseline, a 0.7 mM Mn^2+^ was added.

#### Fura-2 cytosolic Ca ^2+^

After Fura-2AM loading, cells were washed in Ca^2+^-free HBSS (142.6 mM NaCl, 5.6 mM KCl, 5.6 mM glucose, 2.6 mM MgCl_2_, 0.1 mM EGTA, 10 mM HEPES, 0.34 mM Na_2_PO_4_, 4.2 mM NaHCO_3_, 0.44 mM KHPO_4_, pH 7.4). Fluorescence emission was measured at 510 nm after an excitation at both 340 and 380 nm. The ER content was measured using thapsigargin (Tg, 1 μM). The SOCE was measured by adding back 1.8 mM extracellular Ca^2+^.

#### Calcium6 cytosolic Ca ^2+^

After Calcium6 loading, cells were directly recorded and fluorescence emission was acquired at 525 nm after an excitation at 485 nm. 10% FCS in Ca^2+^-free HBSS was added after a 100 s baseline.

### Single cell calcium imaging

100 000 cells plated on 30 mm glass coverslips were loaded with 4 µM Fura-2AM for 45 min. After 3 washes in 1.8 mM Ca^2+^ HBSS, changes in [Ca^2+^]i were monitored using an IX83 (Olympus) inverted microscope-based imaging system equipped with a 40×/1.35 UApo N340 high UV light transmittance oil immersion objective (Olympus), an ORCA-Flash4.0 Digital CMOS camera (Hamamatsu Photonics), a DG-4 Ultra High Speed Wavelength Switcher (Sutter Instruments) and METAFLUOR software (Universal Imaging) for image acquisition and analysis.

### Cell death assays

For cellular viability assessment, eGFP- or eGFP-IP3R3-transfected cells were resuspended in 1 µg/ml Propidium Iodide (PI, 11435392, Fisher Scientific). For mitochondrial membrane potential assay, transfected cells were incubated for 20 min with TMRM (15934294, Fisher Scientific) at 37°C, 5% CO_2_. Cells labeled with either PI or TMRM were processed by flow cytometry (FACS Verse, BD Biosciences). PI or TMRM staining was quantified after gating on GFP-positive cells. Analysis was performed using FlowJo software (BD Biosciences).

### Bioinformatics analyses on publicly available data

Data were extracted from the Firehose legacy TCGA UVM and SKCM databases using cBioPortal (https://www.cbioportal.org) (time stamp: 2023/09/06) or analyzed directly on the cBioPortal platform.

### Statistical analysis

Statistical analysis was made using the GraphPad Prism software (Boston, MA, USA). Normality of the data was determined by a Shapiro-Wilk test then, accordingly, unpaired t-tests were performed for comparison of two groups, or either a one-way or two-ways ANOVA tests. Significance was established when the probability was equal or inferior to 0.05.

## RESULTS

### IP_3_-induced Ca^2+^ signaling is impaired in GNAQ/11^mut^ UVM cells

To investigate the potential remodeling of Ca²⁺ signaling in response to continuous IP3 production in GNAQ/11^mut^ UVM cells, we evaluated Ca^2+^ homeostasis across a panel of melanoma cells. We compared GNAQ/11^Q209L^ UVM cell lines (92.1, MM66 and MP41) to GNAQ/11^Wt^ cells originating either from UVM (OCM-1) or SKCM (M229 and M238) tumors (Figure 1 and Table 1). Initially, we assessed resting Ca²⁺ levels and observed equivalent baselines regardless of mutational status (Figure S1A). Meanwhile, manganese (Mn^2+^) quench assays revealed globally comparable constitutive entries *via* Ca^2+^ conducting channels across all cell lines tested except in MM66 cells where it is smaller (Figure 1A and B). Overall, none of the GNAQ/11^Q209L^ UVM cells tested revealed enhanced resting Ca^2+^ level compared to GNAQ/11^Wt^ cells.

**Figure 1.**
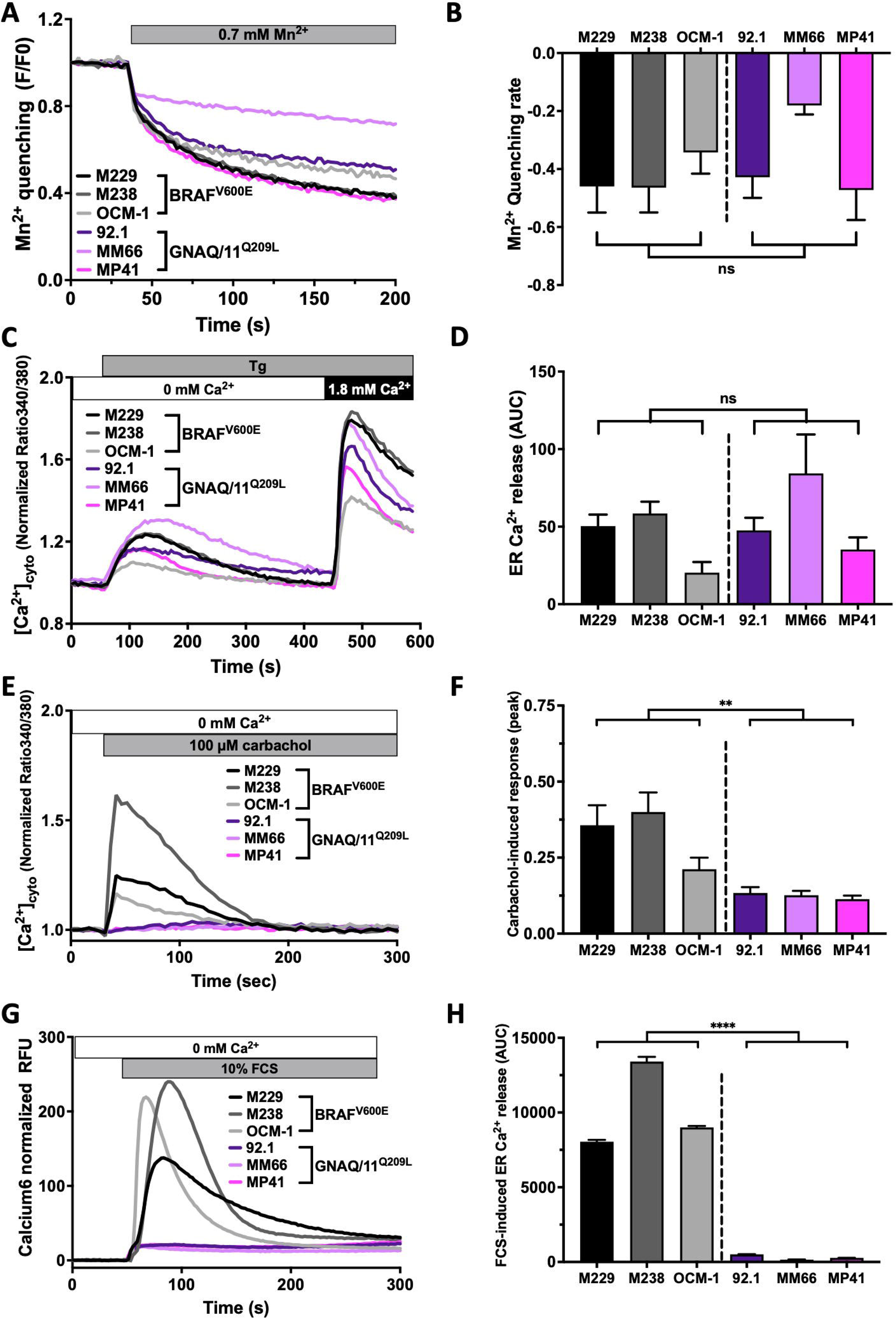
Ca ^2+^ signaling in GNAQ/11 ^Q209L^ UVM cells: resting Ca ^2+^, ER-Ca^2+^ depletion and SOCE. **A-B.** Constitutive Ca^2+^ entry measured by Fura2-based manganese (Mn^2+^) quenching assay in BRAF^V600E^ cell lines (SKCM: M229 and M238, and UVM: OCM-1) compared to GNAQ/11^Q209L^ UVM cell lines (92.1, MM66 and MP41). (A) Representative traces of normalized Fura2 quench (F360), and (B) average Mn^2+^ quenching rate (initial decline, RFU/s) after addition of 0.7 mM Mn^2+^. **C.** Representative traces of normalized thapsigargin (Tg)-induced ER Ca^2+^depletion followed by store-operated Ca^2+^ entry (SOCE) in Fura2-loaded BRAF^V600E^ cell lines compared to GNAQ/11^Q209L^ UVM cell lines. 1 µM Tg was used to deplete ER Ca^2+^ stores in absence of extracellular Ca^2+^. **D.** Average Tg-induced response (area under the curve, AUC) was used to quantify Ca^2+^ release from the ER. **E-F** Representative traces (E) and quantification (F) of the depletion of the ER-Ca^2+^ store induced upon Carbachol stimulation in absence of extracellular Ca^2+^. **G-H** ER-Ca^2+^ store depletion induced upon fetal calf serum (FCS) stimulation in absence of extracellular Ca^2+^. (G) Representative traces of cells loaded with Calcium 6 ((F-F0)/F0). (H) Average FCS-induced Ca^2+^ response (AUC) was used to quantify serum-mediated Ca^2+^ store depletion. Results of at least three independent experiments are shown as mean ± SEM and were analyzed using one-way ANOVA statistical tests, not significant: ns>0.05, ****P<0.0001.

Since the ER is a major intracellular reservoir of Ca^2+^ and a direct target of IP3 signaling, we evaluated whether the steady production of IP3 in GNAQ/11^Q209L^ UVM cells could alter ER-Ca^2+^ content. To do so, we depleted Ca^2+^ from the ER lumen following sarco-ER Ca^2+^-ATPase (SERCA) inhibition by thapsigargin (Tg), in absence of extracellular Ca^2+^ (Figure 1C and S1B). Despite slightly variable rates, GNA11^Q209L^ UVM cells did not display any defective Tg-elicited response, but rather exhibited substantial ER-Ca^2+^ release. If anything, only the GNAQ/11^Wt^ OCM-1 cells exhibited a marginally attenuated depletion. Since ER-Ca^2+^ depletion is directly coupled to SOCE, we next assessed SOCE functionality upon restoration of extracellular Ca^2+^ after ER depletion. Regardless of the mutational status, a solid Ca^2+^uptake was evoked (Figure 1C, D and S1C), indicating that the molecular components of the SOCE are present and fully functional.

Given that Tg bypasses IP3R to induce Ca^2+^ store release (21), we sought to induce ER-Ca^2+^ store depletion *via* IP3 production. In absence of extracellular Ca^2+^, we exposed the cells to carbachol, an agonist of Gα_q/11_-coupled muscarinic acetylcholine receptors (mAChRs) (Figure 1E and F). In that context, GPCRs stimulation straightforwardly induced ER-Ca^2+^ depletion in GNAQ/11^Wt^ cells, while totally inefficient in GNAQ/11^Q209L^ UVM cell lines. Since one could argue that PLCβ activity is already maxed out in UVM cells wherein Gα_q/11_ is constitutively activated, we additionally examined ER-Ca^2+^ depletion upon stimulation with serum, containing a plethora of growth factors, thereby activating a wide variety of surface receptors, including receptor tyrosine kinase (RTK) coupled to PLCγ isozymes. Similarly, serum elicited a significant store depletion resulting in a cytosolic Ca^2+^ wave in GNAQ/11^Wt^ cell lines, whereas in stark contrast, GNAQ/11^Q209L^ UVM cells had completely lost the ability to respond to this IP3-dependent alternative pathway (Figure 1G and H). Collectively, these results indicate that while fully efficient at maintaining Ca^2+^ homeostasis, GNAQ/11^Q209L^UVM cells have implemented a decoupling mechanism between IP3 production and ER-Ca^2+^ store depletion.

### MP41 UVM cells display spontaneous Ca^2+^ oscillations

To further delineate the specificities of Ca^2+^ signaling in GNAQ/11^mut^ UVM cells, and most notably their capacity to withstand a continuous production of IP3, we performed single cell Ca^2+^ imaging experiments. Surprisingly, among the GNAQ/11^Q209L^ UVM cells, the GNA11^mut^ MP41 cell line displayed a unique oscillatory behavior (Figure 2). In presence of extracellular Ca^2+^, long lasting Ca^2+^ oscillations spontaneously arose in the vast majority of GNA11^mut^ MP41 cells (Figure 2A, B and SMovie). Upon extracellular Ca^2+^ removal, oscillations were completely lost, but fully recovered after reimplementing extracellular Ca^2+^, indicating that Ca^2+^ influx is a prerequisite of GNA11^mut^ MP41 cytosolic Ca^2+^ oscillation. Next, we evaluated GNA11 dependency upon *GNA11* depletion with specifically targeting siRNA. In over 95% of the *GNA11-*silenced GNA11^mut^ MP41 cells, Ca^2+^ oscillations were markedly abrogated (Figure 2C and D). Meanwhile, depletion efficiency was validated by the reduction of Gα_11_ expression, and the subsequent diminution of *GNA11* oncogenic signaling using ERK phosphorylation as readout (Figure 2E and S2). Furthermore, Mn^2+^-quench experiments showed that in GNA11^mut^ MP41, *GNA11* depletion significantly impaired basal Ca^2+^ entries (Figure 2F). Hence, our findings revealed that the constitutive activation of the Gα signaling controls specific plasma membrane Ca^2+^ influxes supporting robust Ca^2+^ oscillations in GNA11^mut^ MP41 cells.

**Figure 2.**
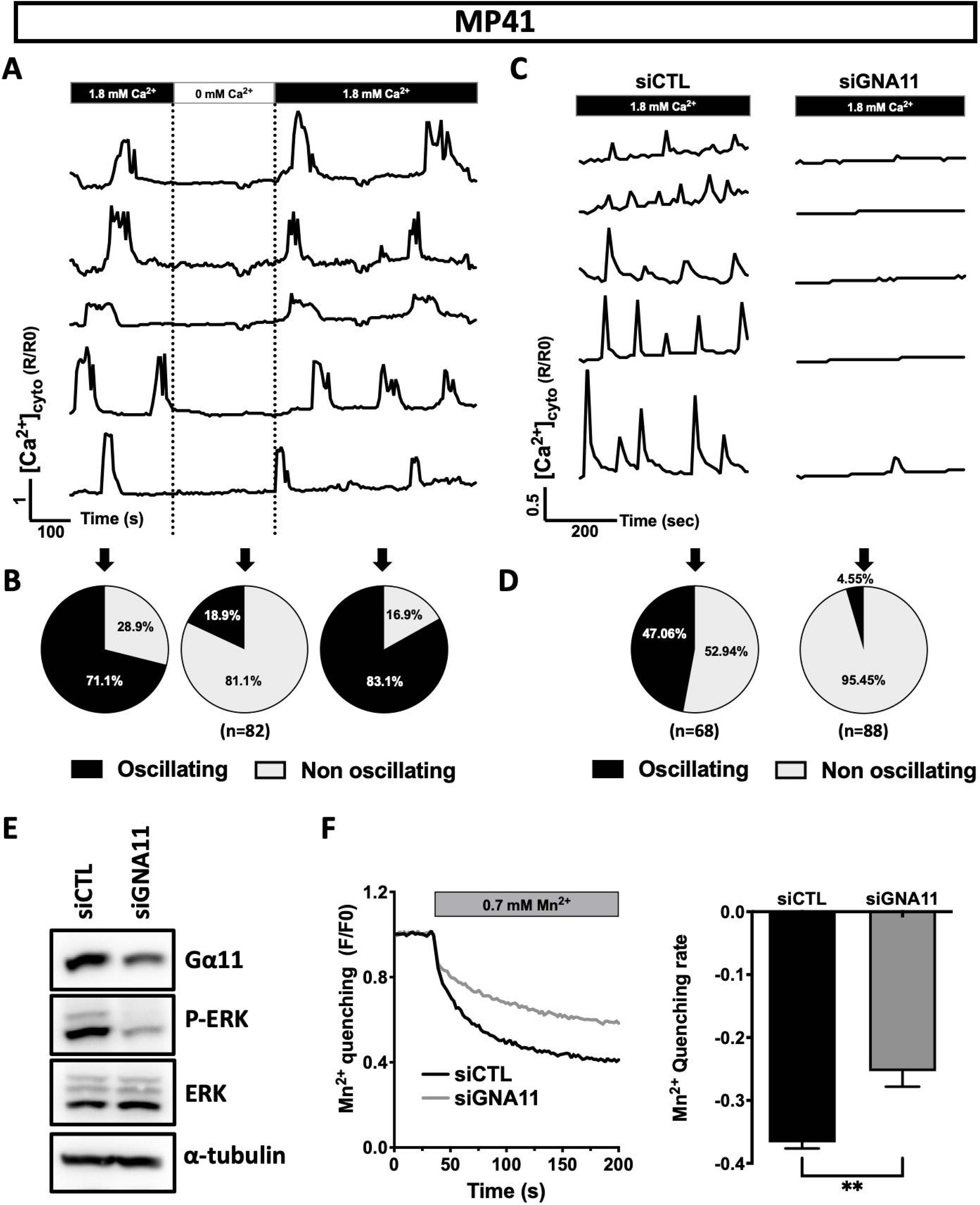
GNA11-dependent spontaneous Ca^2+^ oscillations in the MP41 UVM cell line. **A-B** Fura2-based intracellular Ca^2+^ imaging in MP41 UVM cell line revealing spontaneous Ca^2+^ oscillations, disappearing upon extracellular Ca^2+^ removal and re-occurring upon 1.8 mM extracellular Ca^2+^ addition. (A) Traces of five representative cells are shown. (B) Quantification of non-oscillating (white) and oscillating (black) cells are represented as pie charts for every phase of the experiment (n=82). **C-D** Effect of *GNA11*-targeted siRNA (siGNA11) on spontaneous Ca^2+^ oscillations. (C) Traces of 5 representative cells transfected with either a control siRNA (siCTL) or siGNA11. (D) Quantification of non-oscillating (white) and oscillating (black) cells are represented as pie charts for siCTL (n=68) and siGNA11 (n=88). **E** Effect of GNA11-targeted siRNA (siGNA11) or control siRNA (siCTL) on Gα11, ERK and p-ERK expression assessed by western-blot. α-tubulin was used as a loading control. **F** Effect of siGNA11 on the constitutive Ca^2+^ entry as measured by the Mn^2+^ quenching assay. (Left) Representative traces of normalized Fura2 quench (F360) from siCTL- and siGNA11-expressing cells, and (Right) average Mn^2+^ quenching rate (initial decline, RFU/s) after addition of 0.7 mM Mn^2+^ from four independent experiments (mean ± SEM, unpaired t-test, **P=0.0053).

### Fine-tuned IP3R expression levels in UVM

Because not all GNAQ/11^mut^ UVM cell lines generated spontaneous Ca^2+^ oscillations, we sought to investigate the alternative molecular mechanisms protecting non-oscillatory GNAQ/11^mut^ cells from a constitutive IP3/Ca^2+^ signal. Because IP3 binds to IP3R to mobilize Ca^2+^, we initially analyzed the expression of IP3R transcripts (*ITPR*) (Figure 3A). Despite variabilities among melanoma cell lines, type 1 IP3R (*ITPR*1) remained expressed in GNAQ/11^Q209L^ UVM cells with comparable levels to GNAQ/11^Wt^ SKCM cells, though lower than those detected in GNAQ/11^Wt^ OCM-1 cells. Meanwhile, type 2 (*ITPR*2) and type 3 (*ITPR*3) IP3R expression was significantly reduced in the GNAQ/11^Q209L^ cell lines. Alternatively, the down-regulation of *ITPR* in GNAQ/11^mut^ UVM cells, as compared to GNAQ/11^Wt^ cells, was confirmed on an independent panel of human melanoma cell lines (GSE93666 dataset), wherein *RASGRP3* upregulation in GNAQ/11^mut^ UVM cells supported the robustness of the analysis (Figure S3A) (8). To evaluate the relevance of *ITPR* regulation in patient tumors, we extended our transcriptomic analysis to The Cancer Genome Atlas (TCGA) human melanoma tumor dataset (Figure 3B). To better match the oncogenic mutations and genomic hallmarks characterizing the cell lines used in the present study (*BRAF^mut^* SKCM cells and *GNAQ/11 ^mut^-BAP1^high^* UVM cells) (17–20, 22), we first narrowed down our analysis by retaining only *BRAF^mut^*tumors in the SKCM cohort, and by excluding *BAP1*^low^ tumors from the UVM dataset. Consistent with cell lines, UVM tumors bearing mutations on the GNAQ/11 pathway (n=39) expressed significantly reduced levels of the *ITPR* isoforms as compared to *GNAQ/11^Wt^* tumors (n=123). Importantly, wider analyses performed on the complete SKCM (n=287) and UVM (n=80) TCGA cohorts showed comparable results (Figure S3B).

**Figure 3.**
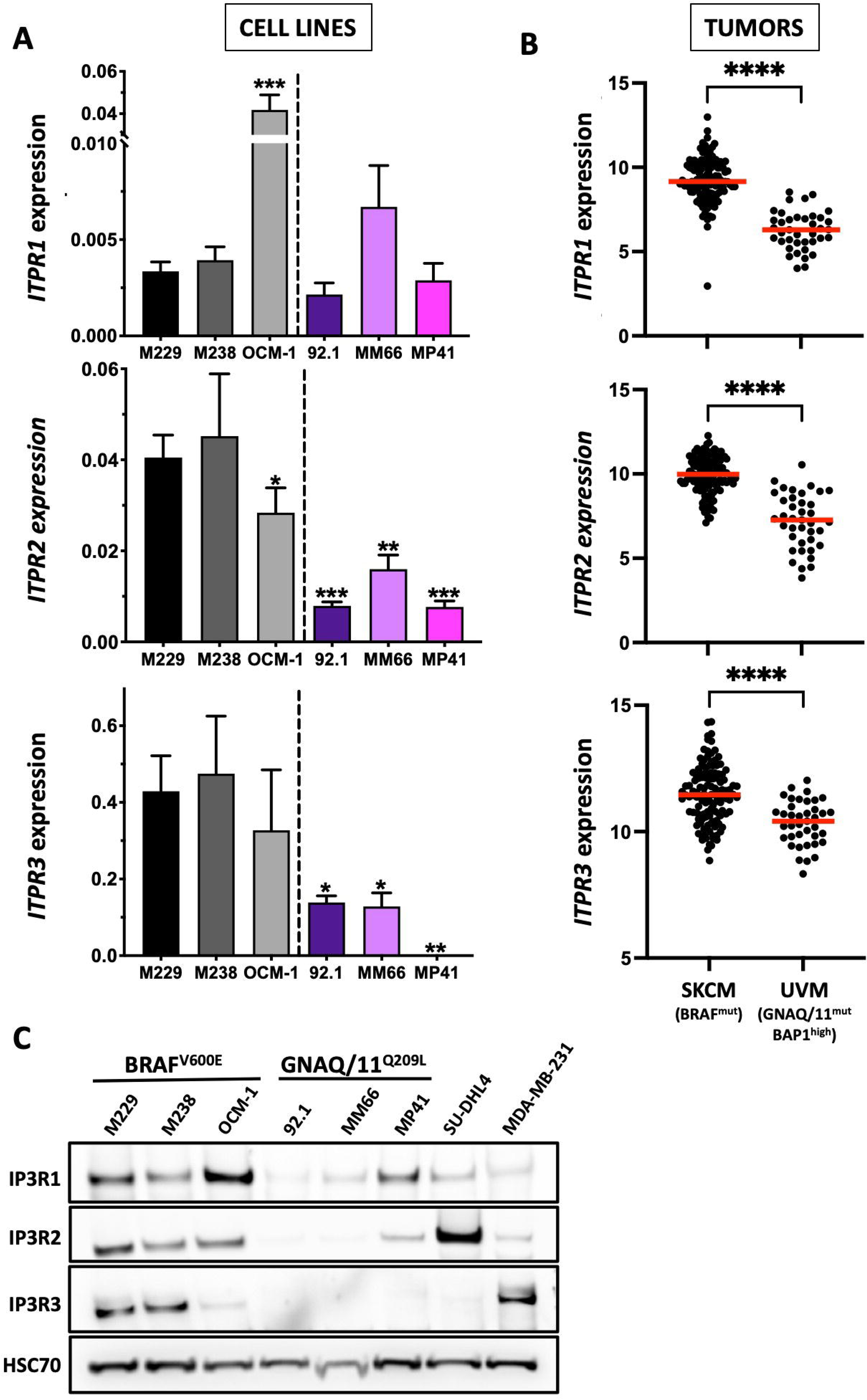
IP3Rs expression in UVM. **A.** Comparison of *ITPR1-3* mRNA expression (ΔΔCt) in BRAF^V600E^ cells (SKCM: M229 and M238, and UVM: OCM-1) *versus* GNAQ/11^Q209L^ UVM cell lines (92.1, MM66 and MP41). qPCR results of at least three independent experiments are represented as mean ± SEM. One-way ANOVA tests were performed, statistical differences are shown for *ITPR1* with OCM-1 cells used as a reference, while for *ITPR2* and *ITPR3,* M238 cells were used as a reference, *P<0.05, **P<0.01, ***P<0.001. **B .** Comparison of *ITPR1-3* mRNA expression (Log2 RNAseq V2 RSEM) levels in BRAF^mut^ SKCM tumors (n=123) and GNAQ/11^mut^/BAP1^high^ UVM tumors (n=39) from the TCGA cohorts. Values are plotted for each individual tumor and the red line indicates the median (unpaired t-test, ****P<0.0001). **C.** IP3R1-3 immunoblotting on BRAF^V600E^ cell lines and GNAQ/11^Q209L^ UVM cell lines. The B-cell lymphoma SU-DHL4 cell line and the breast cancer MDA-MB-231 cell line were used as positive controls for IP3R2 and IP3R3 expression, respectively (62, 63). HSC70 was used as a loading control.

Although expression of *ITPR* transcripts was largely repressed in GNAQ/11^mut^ UVM cells and tumors, weak or moderate levels remained measurable. This prompted us to evaluate protein expression of IP3R channels. We found that IP3R expression was drastically reduced in GNAQ/11^Q209L^ UVM cell lines as compared to GNAQ/11^Wt^ cells (Figure 3C and S3D). Specifically, in the non-oscillating GNAQ^mut^ 92.1 and GNA11^mut^ MM66 cells, IP3Rs were mostly undetectable, with IP3R1 being barely detectable. However, the oscillating GNA11^mut^ MP41 cells exhibited a slightly different behavior, with moderate expression of both type 1 and type 2 IP3Rs, which have been previously associated with Ca²⁺ oscillations (23, 24). In contrast, IP3R3 expression—known to be anti-oscillatory and linked to monophasic Ca²⁺ transients (23–27)—was abolished in all three GNAQ/11^mut^ cell lines. Therefore, the robust Ca^2+^ oscillatory pattern observed in the GNA11^mut^ MP41 cells is presumably associated with the functions of IP3R1/2 functions and the absence of IP3R3. Altogether our results revealed that, in response to GNAQ/11^mut^-induced constitutive IP3 production, the expression of both *ITPR* transcripts and IP3R channel proteins is substantially altered in an isotype-specific manner to fine-tune Ca²⁺ signaling.

Alternatively, IP3R channel activity can be regulated through interactions with various accessory proteins, including the anti-apoptotic B-cell lymphoma 2 (Bcl2) protein which prevents IP3 binding and modulates IP3R-mediated Ca^2+^ signaling. In this study, while IP3R expression was globally reduced in GNAQ/11^mut^ UVM, we also found that Bcl2 expression was substantially enhanced in UVM tumors as compared to the SKCM dataset (Figure S3C). Notably, Bcl2 expression was positively correlated with ITPR1 expression in UVM tumors (Figure S3C). Given the previously described ability of Bcl2 to convert sustained Ca^2+^ signals into Ca^2+^ oscillations (28), we propose that Bcl2-mediated regulation of IP3R1 supports the spontaneous Ca^2+^-oscillatory pattern observed in the GNA11^mut^ MP41 UVM cells. Thus, GNAQ/11^mut^ UVM exhibits complex, multilevel regulatory mechanisms that finely adjust both IP3R expression and activity, effectively restraining excessive IP3/Ca^2+^signaling.

### IP3R3 loss increases survival and protects against Ca^2+^-driven cell death

Considering that type 3 IP3R is involved in Ca^2+^ transfer to the mitochondria during apoptosis and that this specific subtype is undetectable in GNAQ/11^Q209L^ cells, we next examined the impact of IP3R3 loss on UVM cell viability and sensitivity to apoptotic stimuli. First, we evaluated the effect of restoring IP3R3 expression on GNAQ^mut^ 92.1 cell death by monitoring propidium iodide (PI) exclusion and the mitochondrial membrane potential (ΔΨm) (Figure 4A and B). IP3R3 overexpression was sufficient to spontaneously induce cell death, at least partially mediated via the mitochondrial apoptotic pathway. To further investigate the role of IP3R3-dependent cell death, GNAQ^mut^ 92.1 UVM cells were exposed to staurosporine (STS), which has been previously validated to provoke Ca^2+^-driven apoptosis through IP3R signaling (29, 30) (Figure 4C, D, and S4). While STS failed to trigger cell death in control cells, IP3R3 overexpression significantly increased sensitivity to STS-induced apoptosis. Approximately 40% of the IP3R3-expressing cells were permeable to PI and exhibited either a loss (15%) or a gain (25%) in ΔΨm. This dual effect on ΔΨm may reflect an early apoptotic onset dependent on the mitochondrial caspase pathway and a late, Bcl2/caspase-independent step previously described in STS-treated melanoma (31).

**Figure 4.**
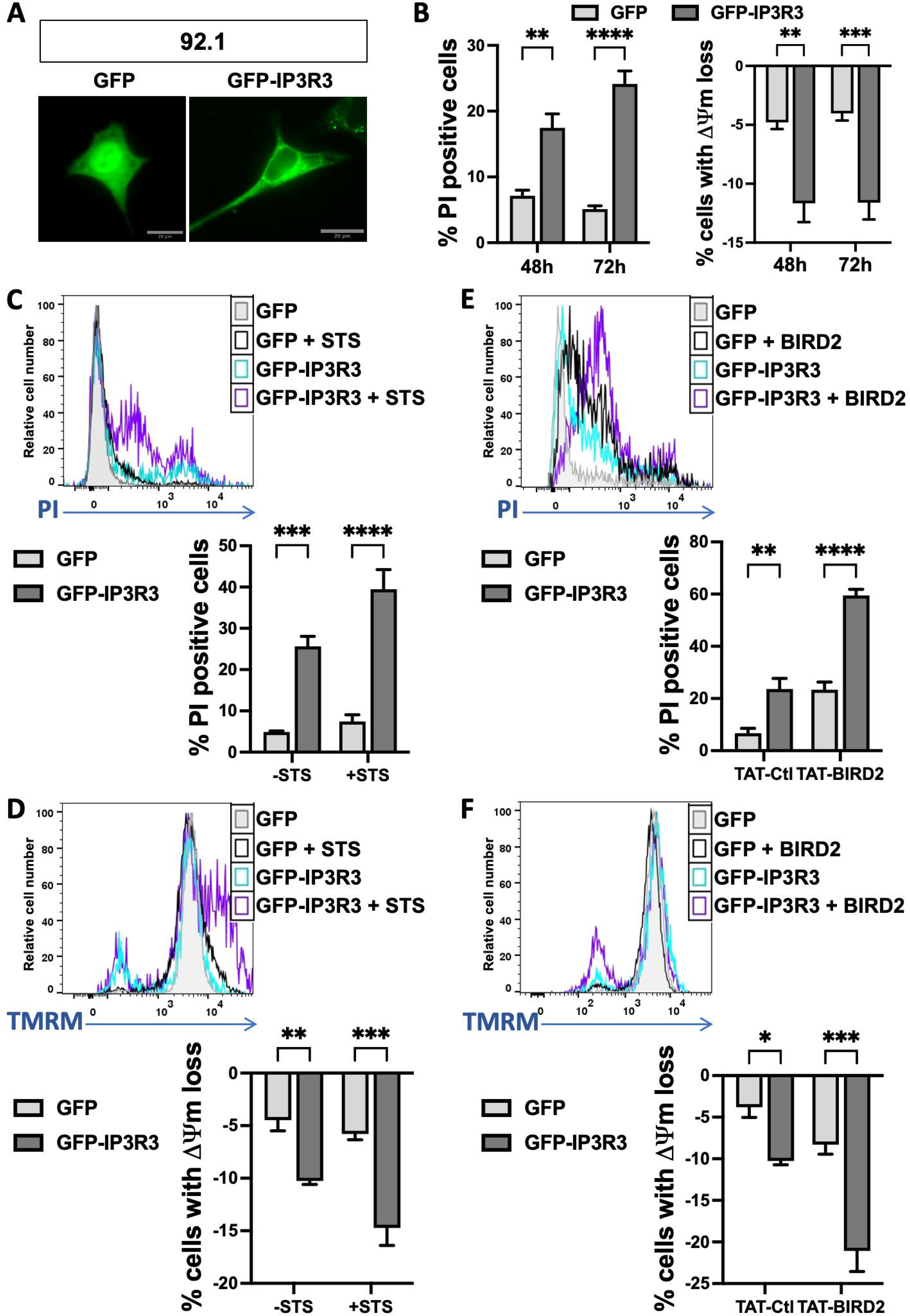
Effect of IP3R3 overexpression on spontaneous cell death and apoptosis induction. **A.** Representative images of 92.1 UVM cells 48 h after transfection with either GFP or GFP-IP3R3. Scale bar, 20 µm. **B.** Cell death was monitored by flow cytometry using both propidium iodide (PI) exclusion and the fluorescent probe TMRM that detects changes in mitochondrial membrane potential (Δψm). Percentages of cells either labeled with PI (left) or that have lost TMRM fluorescence (right) were determined within the GFP positive cell population (to exclude non-transfected cells) 48 and 72 h after transfection. **C-D.** Staurosporine (STS) induced cell death in GFP-IP3R3 but not in control GFP transfected 92.1 cells. Representative flow cytometry histogram (upper panel) and quantification (lower panel) of PI fluorescence (**C**) or TMRM loss of Δψm (**D**) in both GFP and GFP-IP3R3 transfected cells. STS treatment was applied at 0.5 µM for 4 h. **E-F.**TAT-BIRD2 peptide (BIRD2) induced cell death in GFP-IP3R3 transfected 92.1 cells. Representative flow cytometry histogram (upper panel) and quantification (lower panel) of PI fluorescence (**C**) or TMRM loss of Δψm (**D**) in both GFP and GFP-IP3R3 transfected cells. Treatments with either a control (TAT-Ctl) or BIRD2 (TAT-BIRD2) peptide were applied at 10 µM for 2 h. Bar charts are presenting the results of at least three independent experiments as mean ± SEM, with 2-way ANOVA statistical tests, *P<0.05, **P<0.005, ***P<0.001, ****P<0.0001.

To emphasize the role of IP3R3 loss in the protection mechanism developed by GNAQ/11^Q209L^-mutant UVM cells against IP3/Ca^2+^-dependent cell death, we exposed these UVM cells to a decoy peptide that binds to Bcl2. This peptide, Bcl2/IP3R disruptor-2 (BIRD2), effectively blocks Bcl2-IP3R interaction and supports IP3/Ca^2+^-mediated apoptosis (32–34). In IP3R3-overexpressing UVM cells, BIRD2 massively induced apoptosis coupled with a disrupted ΔΨm, while having a more moderate effect on control cells (Figure 4E and F). Taken together, our data demonstrate that when IP3R3 expression is restored, GNAQ/11^Q209L^-mutant UVM cells regain sensitivity to spontaneous cell death as well as to Ca^2+^-driven apoptosis through IP3 signaling.

### GNAQ/11 oncogene-dependent IP3R repression

Next, we sought to explore the mechanisms underlying IP3R repression in GNAQ/11^mut^ UVM cells. Unlike SKCM, UVM represents genetically simple tumors with few copy number alterations (CNAs) and a low mutation burden (35, 36). Analysis of the UVM_TCGA cohort showed that somatic mutations in ITPR genes were very rare, failing to explain the low expression of these genes (Figure S5).

Large genomic studies have subdivided UVM tumors into four molecular and clinical subsets based on somatic mutations and recurrent chromosomal aberrations (36). Good-prognosis class A tumors and intermediate-risk class B tumors retain both copies of chromosome 3 and often show 6p gain. In contrast, high-risk classes C-D tumors frequently exhibit monosomy 3 followed by BAP1 alterations. Gain of 8q is characteristic of aggressive tumors, with higher amplifications in class D tumors. ITPR1 is located on chromosome 3, near BAP1 (cytogenetic bands 3p26.1 and 3p21.1, respectively), while *ITPR3* is found on chromosome 6 (Figure 5A). Given frequent alterations in these regions in UVM, we questioned if CNAs might affect receptor expression. GISTIC prediction in UVM tumors identified frequent shallow deletions of the *ITPR1* and *BAP1* genes that are mostly associated with the monosomy 3 (Figure 5B). Recurrent copy number gains of the *ITPR3* gene related to 6p amplification were also revealed. However, unlike *BAP1*, *ITPR1* and *ITPR3* mRNA expression levels were mostly unaffected by these CNA (Figure 5C). Regarding the *ITPR2* gene found on chromosome 12, despite no CNA detected in UVM tumors, its expression unexpectedly appeared higher in monosomy 3/8q gain poor-prognosis tumors (Figure 5B-C) and negatively correlated with *BAP1* mRNA expression (Figure 5D). This might be due to modifications of transcriptional regulators linked to the genomic abnormalities or to the changes in the global DNA methylation profile observed in *BAP1*-aberrant UVM (36). In any event, our analyses showed that *ITPR* expression is not directly affected by recurrent mutations or CNAs in UVM.

**Figure 5.**
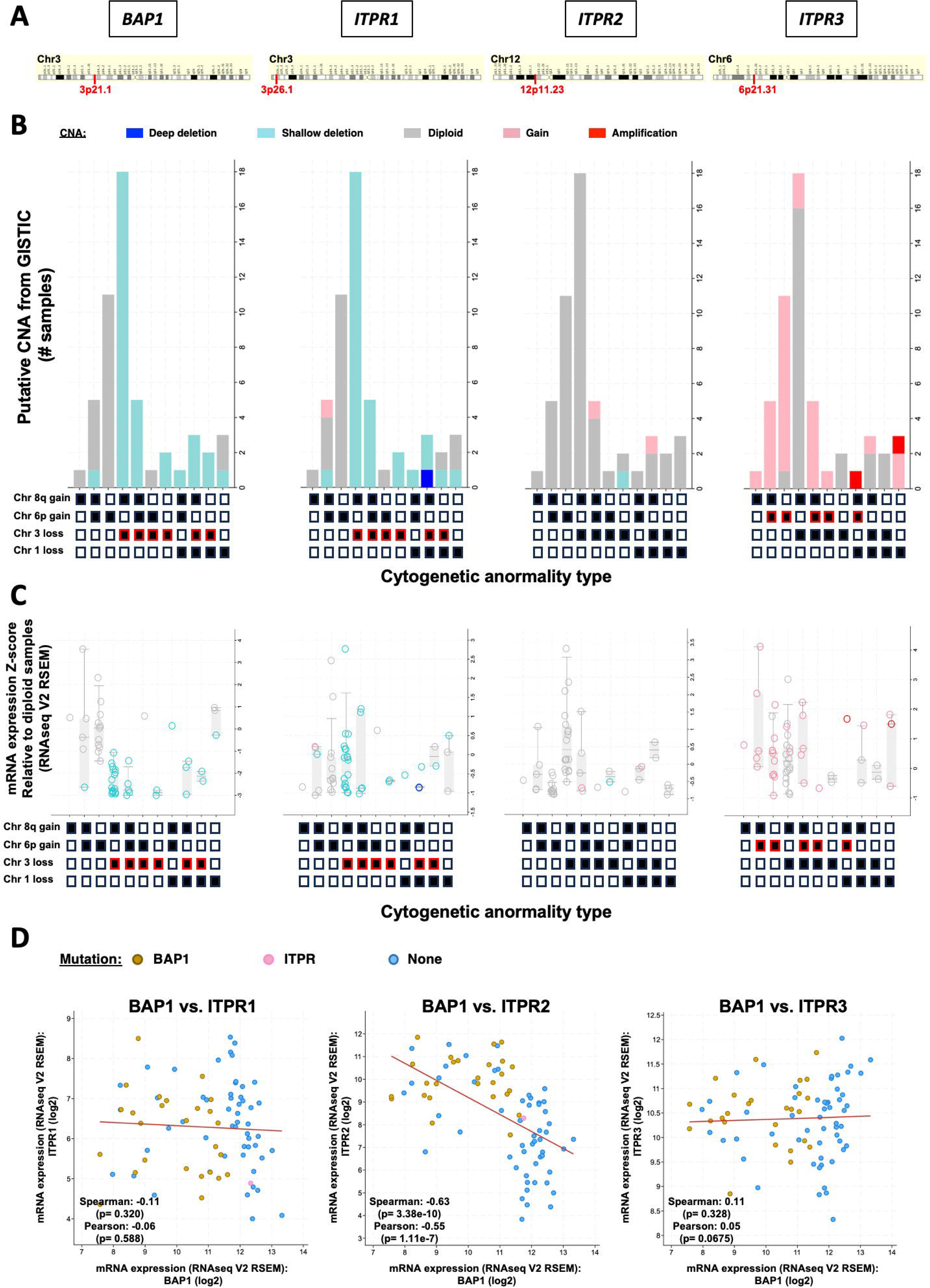
Impact of UVM cytogenetic abnormalities and CNA on ITPRs expression. **A.** Human genomic location of *BAP1* and *ITPR1-3* genes with specific cytogenetic bands in red. **B-C.** In the UVM_TCGA cohort (retrieved from https://www.cbioportal.org), for each type of frequent UVM cytogenetic abnormalities (highlighted in red when the aberration occurs on the same chromosome as the gene of interest), the number of samples/patients with putative GISTIC-detected copy number alteration (CNA) (**B**) or the mRNA expression Z-score relative to diploid samples for each gene (**C**) are plotted. Types of alterations are color-coded on the basis of the legend. 52 samples from the UVM_TCGA cohort with available data in both profiles (axes) are presented. **D.** In the same UVM cohort, BAP1 expression is negatively correlated with *ITPR2,* but not with *ITPR1* or *ITPR3* expression.

As IP3R repression is specific to GNAQ/11^mut^ UVM but not GNAQ/11^wt^ cells, we assessed whether the oncogenic Gαq/11 signaling pathway controls IP3R repression (Figure 6A). We first interrogated RNAseq data issued from short-term cultures of UVM biopsies (37) that were submitted *ex-vivo* to Gα_q/11_ inhibition with FR900359 (Figure 6B). This revealed enhanced expression of at least one ITPR isotype within each treated tumor (Figure 6B), implicating constitutive Gαq/11 activation in IP3R modulation. We further substantiated the role of the Gα_q/11_ oncogenic pathway in regulating *ITPR* expression by directly inhibiting Gα_q/11_ with YM-254890 in UVM cell lines (Figure 6C, D and S6). Gα_q/11_ inhibition restored *ITPR*/IP3R expression in GNAQ^mut^ 92.1 cells, but not in GNAQ/11^Wt^ OCM-1 cells, while the downstream phosphorylation of ERK was efficiently inhibited in both cell lines. Gα_q/11_ direct inhibition is therefore sufficient to disengage the mechanism hindering *ITPR*/IP3R expression, revealing that Gα_q/11_ signaling pathway actively controls IP3R repression in UVM.

**Figure 6.**
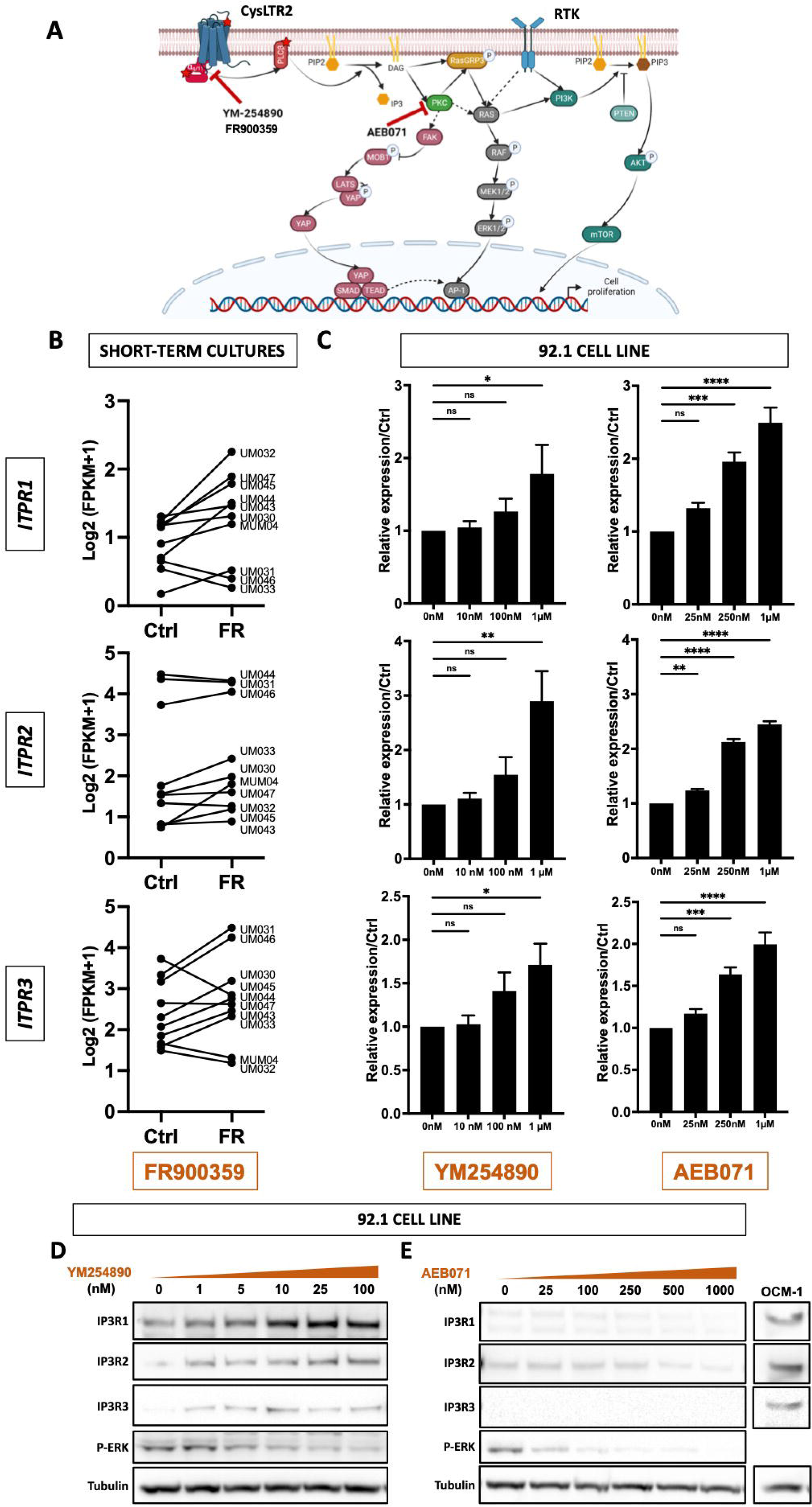
IP3R expression dependency towards the oncogenic Gα_q/11_ signaling pathway. **A.** Schematic representation of the main pathways involved in UVM, with an emphasis on the inhibitors used in this study and their targets. **B.** *ITPR1-3* mRNA expression in short-term culture issued from *GNAQ/11^mut^*UVM biopsies (37) that were submitted *ex vivo* to a treatment with the Gα_q/11_ inhibitor FR900359 (FR) or not (Ctrl). **C.** *ITPR1-3* mRNA expression in the *GNAQ/11^mut^* 92.1 cell line, 24 h after treatment with the indicated concentration of either the Gα_q/11_ inhibitor, YM-254890 or the pan-PKC inhibitor, AEB071 relative to the control condition (vehicle). qPCR results (2^ΔΔCt^ analysis) of three independent experiments are represented as mean ± SEM (one-way ANOVA, *P<0.05, **P<0.005, ***P<0.001, ****P<0.0001). **D.** Immunoblots of IP3R channels expression in 92.1 *GNAQ/11^mut^* cell line 24 h after treatment with the indicated concentration of either the Gα_q/11_ inhibitor, YM-254890 or the pan-PKC inhibitor, AEB071. OCM-1 lysates are used as positive control of IP3R1-3 expression.

One of the first known targets following Gα_q/11_ activation is the DAG-activated PKC, controlling the RasGRP3/MAPK signaling branch that drives UVM uncontrolled proliferation (8) (Figure 6A). Hence PKC inhibition has been actively investigated and evaluated in clinic, but giving unsatisfying responses (38). To determine whether the PKC/RasGRP3/MAPK signaling branch was involved in IP3R repression, we assessed *ITPR* expression following treatment with sotrastaurin (AEB071), a pan-PKC inhibitor used in clinical trials (39). Upon PKC inhibition, *ITPR* expression was markedly enhanced in GNAQ^mut^ 92.1 cells, while unaffected in GNAQ/11^Wt^ OCM-1 cells (Figure 6C and S6). However, despite increased transcripts, IP3R proteins remained barely detectable (Figure 6D). Unlike Gαq/11 inhibition, PKC inhibition does not reduce IP3 production, potentially leading to IP3R degradation via overstimulation (40). Together, these results demonstrate that while mutations and CNAs do not drive IP3R repression, the oncogenic Gαq/11/DAG/PKC/RasGRP3/MAPK signaling pathway actively suppresses IP3R expression.

## DISCUSSION

The main finding of this study is that, while preserving Ca^2+^ homeostasis to ensure survival, GNAQ/11^mut^ UVM evade IP3/Ca^2+^-driven cell death by altering the expression and activity of IP3R channels. Although the sustained production of IP3 in GNAQ/11-mutant UVM cells has been clearly demonstrated recently (41), its effect on Ca^2+^ signaling has been disregarded. This study underscores the need for a detailed understanding of IP3R-dependent protective mechanisms to improve therapeutic strategies for UVM patients.

### Finely-tuned Ca ^2+^ signaling in UVM to simultaneously preserve survival and elude cell death

Cells typically use transient elevations of cytosolic Ca^2+^ to provide finely tuned survival signals while avoiding prolonged Ca^2+^ rises that can activate proteases, induce ER stress, and cause mitochondrial Ca^2+^ overloading. In this study, we demonstrated that GNAQ/11^mut^ UVM cells have effectively decoupled PLC-mediated IP3 generation from ER-Ca^2+^ depletion, without disrupting resting Ca^2+^ concentration, intracellular ER-Ca^2+^ loading, or the SOCE machinery. However, as the primary intracellular Ca^2+^ storage organelle, the ER requires tightly regulated Ca^2+^ concentrations for proper protein synthesis and folding. To stabilize both cytosolic and ER Ca^2+^ concentrations, ER-Ca^2+^ release through IP3R channels, along with the resulting Ca^2+^ influx mediated by the highly sensitive STIM2/Orai1 machinery, is essential (42). Consequently, GNAQ/11^mut^ UVM cells must preserve a regulatory loop to maintain intracellular Ca^2+^ homeostasis and prevent ER stress. While we attribute the IP3/Ca^2+^ uncoupling process to the downregulation of IP3R expression, the remaining minimal expression of IP3R1 may be sufficient to sustain steady-state ER-Ca^2+^ content and support Ca^2+^ transfer to mitochondria.

IP3R-mediated Ca^2+^ release from the ER can “quasi-synaptically” transfer into the mitochondria *via* the so-called mitochondria-associated ER membranes (MAM). This coupling that permeates Ca^2+^ across the mitochondrial outer membrane occurs *via* the IP3R and voltage-dependent anion-selective channel protein 1 (VDAC1) (43, 44). Mitochondria can therefore buffer substantial amounts of Ca^2+^ and influence cytosolic Ca^2+^ changes. Moreover, Ca^2+^-release events from the ER mediated by IP3R channels are involved in sustaining mitochondrial bioenergetics (45). Indeed, the sequestration of Ca^2+^ by mitochondria and the subsequent increase in respiration is crucial for coupling cellular activity with energy demand. Conversely, excessive mitochondrial Ca^2+^ uptake, notably through IP3R3, leads to the activation of the intrinsic apoptotic pathway *via* release of pro-apoptotic factors (46). Similar to ER-Ca^2+^, IP3R-mediated mitochondrial Ca^2+^ can thereby support both survival and pro-apoptotic cellular functions, according to the magnitude of Ca^2+^ transfer, corroborating the tight spatiotemporal control of IP3R levels in UVM.

Meanwhile, IP3R knock-out studies have shown that, at least in Hela cells, IP3R are dispensable for maintaining cells metabolically active and alive (47). Hence, by suppressing IP3R expression, and thus abrogating IP3R-mediated ER-Ca^2+^ release, GNAQ/11^mut^ UVM escape ER stress, mitochondrial overload and ultimately cell death. Accordingly, our work demonstrates that IP3R3 overexpression in GNAQ/11^mut^ UVM cells not only constitutively induced cell death, but also reversed their resistance to pro-apoptotic inducers, such as STS or BIRD2. Whether IP3R3-overexpressing GNAQ/11^mut^ UVM “spontaneous” cell death could be impeded by inhibiting GNAQ/11^mut^-induced constitutive production of IP3 would be a relevant analysis to conduct. As a matter of fact, Gα_q/11_ inhibition with FR900359 has been shown to be hardly efficient at inducing cell death, presumably because the constitutive production of IP3 is *de facto* abrogated (48). Another important parameter influencing ER-mitochondrial Ca^2+^ transfer, and thereby apoptosis, is the regulation of IP3R activity by modulatory factors. Notably, the anti-apoptotic protein, Bcl2, which as an inhibitor of the IP3R channel activity greatly impact Ca^2+^ dynamics and prevent pro-apoptotic Ca^2+^ release (49, 50). Accordingly, in IP3R3 overexpressing UVM cells, BIRD2 induced a massive loss of mitochondrial membrane potential implying mitochondrial Ca^2+^ overload that triggered cell death. Hence, we argue that the tight regulation of IP3R availability and activity represents a pivotal node to control Ca^2+^ signaling and protect *GNAQ/11^mut^* UVM against IP3/Ca^2+^-driven cell death.

### Oncogenic UVM cells survival and progression do not require SOCE

We showed that GNAQ/11^mut^ UVM cells have abrogated the IP3-dependent release of Ca^2+^ from intracellular stores upon membrane receptor stimulation by growth factors. Yet, it is well established that IP3-mediated Ca^2+^ depletion of the ER triggers the activation of SOCE. Using recruitment of SOCE, growth factor receptors generate subtle, localized and transient intracellular Ca^2+^ waves or spikes which are required for cell proliferation and survival. While numerous reports indicate that SOCE contributes to oncogenesis by promoting malignant behavior, such as tumor growth, angiogenesis and metastasis of tumor cells, including in metastatic UVM cells (51), other evidence has emerged that in specific cellular contexts, such as SKCM, invasion and metastasis are rather promoted by SOCE suppression (52, 53). This would suggest that by suppressing IP3R-mediated SOCE, GNAQ/11^mut^ UVM promotes pro-oncogenic phenotypes, and that the extent to which IP3R tunes SOCE is determined by the strength of Gq signaling. Finally, raising the question of adaptive changes within the SOCE pathway in tumor cells.

### IP3R subtype-specific regulation determine different Ca^2+^-dependent protection mechanisms

It has been well established that with cytosolic Ca^2+^ oscillations, cells not only send out encoded signals to modulate cellular activity, but also avoid deleterious effects of elevated cytosolic Ca^2+^ (23, 28). Cytosolic Ca^2+^ oscillations are IP3R subtype-dependent. Upon a dynamic production of IP3, IP3R1 and 2 crucially contribute in generating these oscillations, whereas IP3R3 functions as an anti-oscillatory unit (24). Here, we evidenced that GNA11^mut^ MP41 UVM cells display spontaneous Ca^2+^ oscillations dependent on both extracellular Ca^2+^ and *GNA11^mut^*-mediated production of IP3. Consistently, in these UVM cells, but not in all GNAQ/11^mut^ UVM cells, IP3R1 and 2 remained expressed whereas IP3R3 was undetected. In addition, our transcriptomic analysis reveals that IP3R1 expression correlates with Bcl2 expression in UVM tumors. Meanwhile, Bcl2 has been described as able to convert sustained IP3-mediated high-amplitude Ca^2+^ signals into Ca^2+^ oscillations (28). Hence, we argue that GNA11^mut^ MP41 UVM cells developed a protective mechanism against long lasting IP3-dependent Ca^2+^ signals through the generation of robust cytosolic Ca^2+^ oscillations involving Bcl2-dependent regulation of IP3R1 and IP3R2 activities. *GNA11*-driven Ca^2+^ oscillations occurring in resting GNA11^mut^ MP41 cells mediate pro-survival functions, notably by preserving Ca^2+^ homeostasis, but may also support tumor aggressive behavior (28, 54, 55). Indeed, Orai/STIM1-mediated Ca^2+^ oscillation signals were reported to promote melanoma invasion (56). A higher number of cell lines should nevertheless be tested to firmly conclude on the protective role of Ca^2+^ oscillations in *GNA11^mut^* UVM. These results however support the idea that IP3R subtype-specific expression acutely shapes Ca^2+^ signaling patterns and its subsequent functions.

### Mechanisms of IP3R Repression in GNAQ/11-Mutant Uveal Melanoma ?

By shutting down cell death sensitivity while simultaneously preserving pro-survival signals, the regulation of IP3R represents a key node at the intersection of cell death and proliferation in the pathogenesis of GNAQ/11-mutant UVM. Although we were unable to fully elucidate the regulatory mechanisms that maintain the delicate balance between repression and expression, we excluded certain mechanisms and highlighted the complexity of IP3R regulation. Notably, our analyses revealed that in UVM tumors, ITPR expression is not directly influenced by recurrent mutations or copy number alterations (CNA).

Nevertheless, we provide evidence of a dependency of ITPR/IP3R repression on the GNAQ/11 oncogenic mutation. Specifically, Gαq/11 inhibition rescued IP3R expression at both transcript and protein levels in GNAQ/11-mutant UVM cell lines, but not in BRAF-mutant UVM cells. Similarly, in short-term cultures derived from UVM patient biopsies, Gαq/11 inhibition promoted ITPR expression. Interestingly, not all ITPR isoforms were equally affected; rather, one or two isoforms were markedly upregulated upon Gαq/11 inhibition. Consistent with this, we observed that in GNAQ/11-mutant UVM cell lines, while overall ITPR levels were significantly reduced, at least one ITPR isoform maintained minimal expression. Further analysis revealed that, in addition to transcriptional repression, IP3R channels are subjected to post-transcriptional and/or translational regulation, limiting their availability. It is important to note that IP3-binding affinity varies depending on the IP3R isoforms, with the proposed order being IP3R2 > IP3R1 > IP3R3 (57). Consequently, alterations in even small amounts of a single isoform could critically impact Ca²⁺ signaling.

Alternatively, we cannot exclude other IP3R regulatory mechanisms, such as modulation by gating regulators including ATP, cAMP, H+ and the redox environment, or covalent modifications, phosphorylation, and ubiquitination (58, 59). Not to forget proteasomal degradation known to target activated IP3R (41, 60) or the huge array of proteins that associate with and regulate IP3R activity and distribution. In that specific context, we highlighted a pivotal role for Bcl2 in regulating IP3R activation in GNAQ/11^mut^ UVM tumors. This was supported by *i*, the magnitude of the apoptotic response induced by inhibiting Bcl2/IP3 interaction, *ii*, the enhanced expression of Bcl2 in UVM tumors, and *iii*, the positive correlation between Bcl2 and IP3R1 expression in UVM tumors. Moreover, Bcl2-mediated modulation of IP3R activity has been reported to convert massive Ca^2+^ elevations that lead to cell death, into lower amplitude, oscillatory Ca^2+^ signals that favor cell survival (28). Altogether, our study reveals that GNAQ/11^mut^ UVM implement complex multi-step checkpoints to tightly maintain IP3R expression and activity within limits, as a critical mechanism to elude IP3/Ca^2+^-triggered cell death while promoting tumorigenesis.

### Towards new combined therapeutic strategies

Finally, given the prevalence of oncogenic aberrations in *GNAQ/11* signaling, agents targeting downstream effectors of the Gα_q/11_ pathway, such as MEK (selumetinib and trametinib) and PKC (sotrastaurin) have been the object of numerous studies and clinical trials (61). However, as with chemotherapeutic approaches, unsatisfying therapeutic outcomes have been reported. In this original study, we demonstrate that in addition to the previously described proliferative functions through the DAG branch, GNAQ/11^mut^signaling pathway also protects against IP3/Ca^2+^-driven cell death *via* a fine control of IP3R channels as a pivotal safety mechanism during UVM tumorigenesis. In practice, direct inhibition of Gα_q/11_ can promote G1 cell cycle arrest, but only induces weak apoptosis (48), presumably because albeit IP3R expression is recovered, IP3 constitutive production is withheld. Thus, although upstream inhibition of the GNAQ/11 signaling pathway efficiently dampened cell proliferation, it appears not sufficient to induce sustained cell death and eliminate UVM cells. This suggests that targeting multiple key nodes of the GNAQ/11 signaling pathway could achieve better therapeutic results. Our study highlights IP3R-evoked Ca^2+^ signal as a key therapeutic node, where the prolonged production of IP3 triggers cell death. Since overstimulated IP3R undergo degradation, as is the case upon PKC inhibition, we speculate that inhibiting IP3R degradation conjointly to PKC activity, for instance, may be an efficient way to maintain IP3/IP3R balance and induce UVM cell death together with a hindered proliferation. Therefore, a wider understanding of the multiple functions of GNAQ/11 signaling promoting survival and simultaneously eluding cell death may ultimately be exploited for the design of novel therapeutics strategies available for UVM patients.

From a broader perspective, by revealing novel mechanistic insights into how GNAQ/11 constitutive activation leads to UVM oncogenesis by promoting survival while simultaneously avoiding IP3/Ca^2+^-driven cell death, our findings open new possibilities for the design of novel combined therapeutics strategies.

## Abbreviations

(BAP1): BRCA1-associated protein 1
(Bcl-2): B-cell lymphoma 2
(Ca^2+^): Calcium ions
(CNA): copy number alteration
(cysteinyl leukotriene receptor 2): CYSLTR2
(DAG): diacylglycerol
(*GNAQ*): guanosine nucleotide-binding protein Q
(*GNA11*): guanosine nucleotide-binding protein alpha-11
(GPCR): G protein-coupled receptor
(IP3): inositol 1,4,5-triphosphate
(*ITPR*) or protein (IP3R): inositol trisphosphate receptor gene
(MAM): mitochondria-associated ER membrane
(PIP2): phosphatidylinositol 4,5-bisphosphate
(PKC): Protein Kinase C
(PLCβ): phospholipase C beta
(RasGRP3): RAS Guanyl Releasing Protein 3
ATPase (SERCA): sarco-ER Ca^2+^
(SF3B1): splicing factor 3B1
(SKCM): skin cutaneous melanoma
(SOCE): store-operated Ca^2+^ entry
(SRSF2): serine and arginine rich splicing factor 2
(STIM): stromal interaction molecule
(TCGA): The Cancer Genome Atlas
(UVM): uveal melanoma

## ACKNOWLEDGMENTS

We would like to thank Dr Lise Rodat Despoix and Dr Olivier Mignen,for their help and critical discussions during the course of our study. We are also grateful to Dr. Samar Alsafadi and Dr. Sergio Raman Roman from the Institut curie translational research department for sharing resources. The transcriptomic analyses shown here are in part based upon data generated by the TCGA Research Network: https://www.cancer.gov/tcga. This work was funded by the Région Nouvelle Aquitaine (Chaire Universitaire Canaux Calciques et Mélanome, A.P, #16/RALPC-P-R-23). C.G was a doctoral fellowship recipient from the Fondation de France (Programme “Allocations jeunes chercheurs en ophtalmologie et neuro-ophtalmologie”; #00099524).

## Author Contributions

Conceptualization, S.L-P, G.B and A.P; Formal analysis, C.G., S.L-P, A.P.; Investigation, C.G., L.R., S.M., S.L-P, A.P.; Project Administration: A.P; Resources, C.B. and S.T-D; Supervision: S.L-P, G.B and A.P; Visualization: C.G, S.L-P and A.P; Writing – Original Draft, S.L-P and A.P; Funding Acquisition, A.P.

All authors have read and acknowledged the content of the manuscript.

