## Supplemental Figures 1-6 for "Oncogenic GNAQ/11-induced remodeling of the IP3/Calcium signaling pathway protects Uveal Melanoma against Calcium-driven cell death"

***Supplementary Figure 1. $Ca^{2+}$ signaling in $GNAQ/11^{Q209L}$ UVM cells: resting $Ca^{2+}$and SOCE.***

**A.** Average resting $Ca^{2+}$ levels in $BRAF^{V600E}$ cells (SKCM: M229 and M238, and UVM: OCM-1) *versus*$GNAQ/11^{Q209L}$ UVM cell lines (92.1, MM66 and MP41) measured with Fura2 (F340/F380). **B-C.** Average SOCE responses in $BRAF^{V600E}$ *versus* $GNAQ/11^{Q209L}$ UVM cell lines in presence of 1.8 mM extracellular $Ca^{2+}$. Initial slopes (B) and peak magnitude (C) were used to quantify $Ca^{2+}$ entry. Results of at least four independent experiments are shown as mean ± SEM, with one-way ANOVA statistical tests, not significant: ns P>0.05.

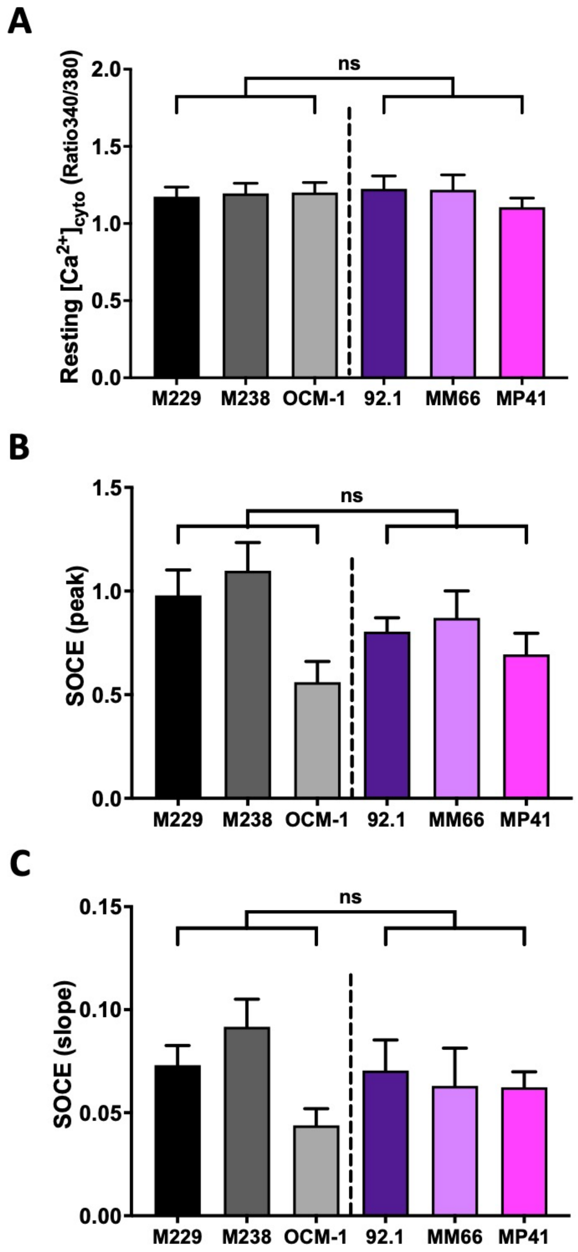

***Supplementary Figure 2. Validation of GNA11 silencing.***

Effect of *GNA11*-targeted siRNA (siGNA11) or its siRNA control (siCTL) on G$\alpha_{11}$, ERK and p-ERK expression. **Bar graph represent the q**uantification of G$\alpha_{11}$ expression (A) and p-ERK/ERK ratio (B) relativized to siCTL. Results of at least six independent experiments are shown as mean ± SEM, with an unpaired t-test, **P<0.01, ****P<0.0001.

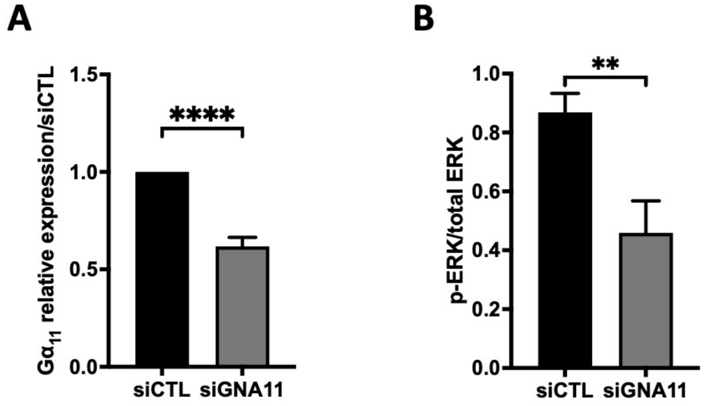

***Supplementary Figure 3. IP3R and Bcl2 expression in UVM cell lines and tumours.***

**A.** Heatmap (expression Z-score analysed from GSE93666 microarray data [8]) comparing *ITPR1-3* and *RasGRP3* expression profiles between five GNAQ/11$^{mut}$ UVM cell lines (GNAQ$^{mut}$: 92-1, MEL202, OMM1.3; GNA11$^{mut}$: UPMD-1, OMM-GN11) and five GNAQ/11 wild-type melanoma cell lines harbouring either BRAF (SK-MEL-5 from SKCM origin and MUM2C from UVM origin) or NRAS mutations (SK-MEL-2, MM415 and MM485, all from cutaneous origins). **B.** Comparison of *ITPR1-3* mRNA expression (Log2 RNAseq V2 RSEM) levels in all SKCM (n=287) *versus* UVM (n=80) tumours from the TCGA cohorts. Values are plotted for each individual tumour and the red line indicates the median. Statistical differences were analysed with an unpaired t-test, ****P<0.0001. **C.** Left panel: comparison of *BCL2* mRNA expression level (Log2 RNAseq V2 RSEM) in SKCM (n=287) *versus* UVM (n=80) tumours from the TCGA cohorts. Values are plotted for each individual tumour and the red line indicates the median. Statistical differences were analysed with an unpaired t-test, ****P<0.0001. Right panel: correlation between *BCL2* and *ITPR1* mRNA expression in UVM tumours. **D.** Quantification of IP3R1-3 protein expression in GNAQ/11$^{Q209L}$ UVM cell lines compared to BRAF$^{V600E}$ cells. Results of at least three independent experiments are represented as mean ± SEM. One-way ANOVA tests were performed, with statistical differences shown for *ITPR1, ITPR2* and *ITPR3, using* M229 cells as a reference, not significant: ns P>0.05, *P<0.05, ****P<0.001.

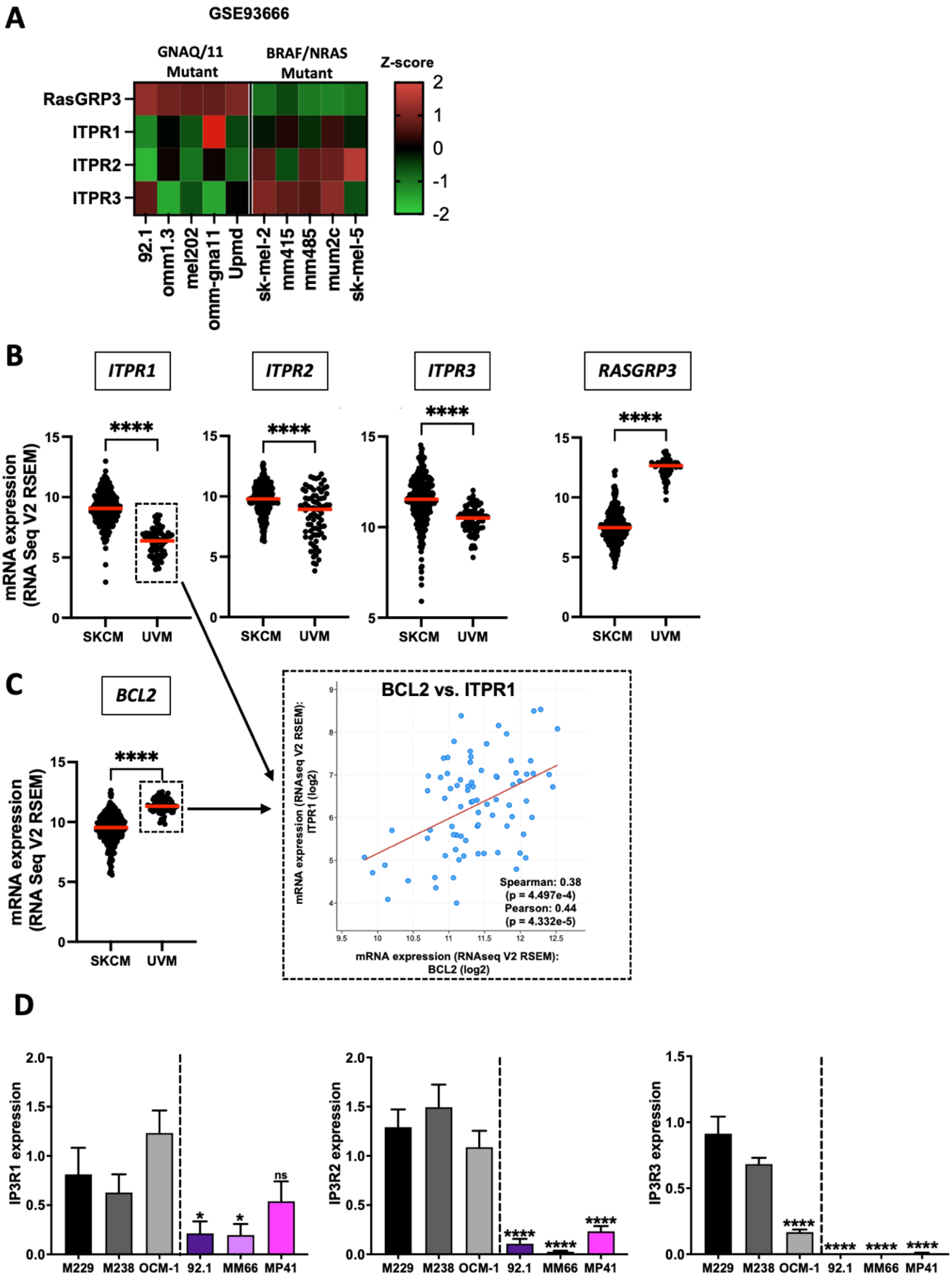
A
GSE93666
GNAQ/11 Mutant
BRAF/NRAS Mutant
Z-score
RasGRP3
ITPR1
ITPR2
ITPR3
92.1
omm1.3
mel202
omm-gna11
Upmd
sk-mel-2
mm415
mm485
mum2c
sk-mel-5
B
ITPR1
ITPR2
ITPR3
RASGRP3
mRNA expression (RNA Seq V2 RSEM)
SKCM
UVM
C
BCL2
BCL2 vs. ITPR1
mRNA expression (RNAseq V2 RSEM): ITPR1 (log2)
mRNA expression (RNAseq V2 RSEM): BCL2 (log2)
Spearman: 0.38
(p = 4.497e-4)
Pearson: 0.44
(p = 4.332e-5)
D
IP3R1 expression
IP3R2 expression
IP3R3 expression
M229
M238
OCM-1
92.1
MM66
MP41
ns

***Supplementary Figure 4. STS and BIRD2 effect on mitochondrial membrane potential.***

Quantification of the gain of mitochondrial membrane potential (Δψm) in GFP- and GFP-IP3R3 transfected 92.1 UVM cell line, after treatment with either staurosporine (STS) (left panel) or BIRD2 peptide (right panel). Δψm was monitored by flow cytometry using the TMRM fluorescent probe. Results of three independent experiments are represented as mean ± SEM, with a two-way ANOVA test, not significant: ns P>0.05, **P<0.01, ***P<0.001.

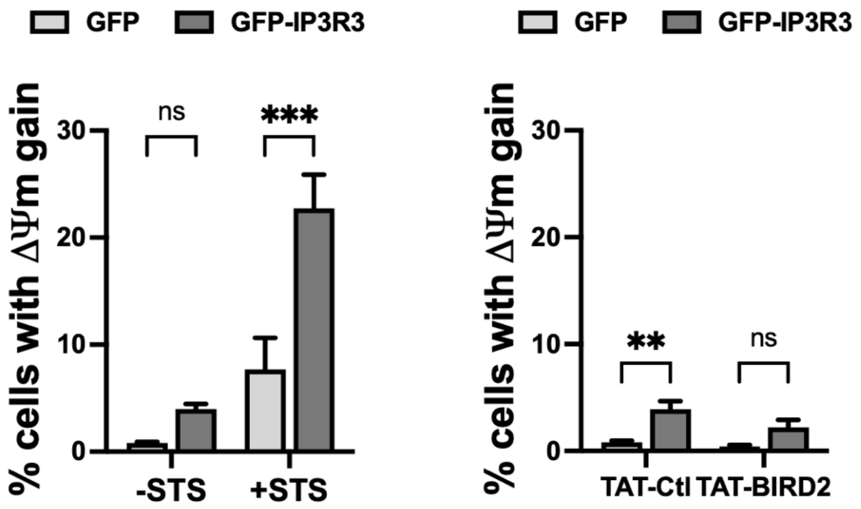

***Supplementary Figure 5. ITPR genes genetic alterations in UVM.***

The oncoprint analysis presents data obtained from c-BioPortal (time stamp: 2023/09/08) by querying the 80 samples/patients of the Uveal Melanoma (TCGA, Firehose Legacy) database. Types of alterations are color-coded according to the legend. The indicated percentages represent the alteration frequency in the specified gene. Each column represents a patient/sample and unaltered cases were not included.

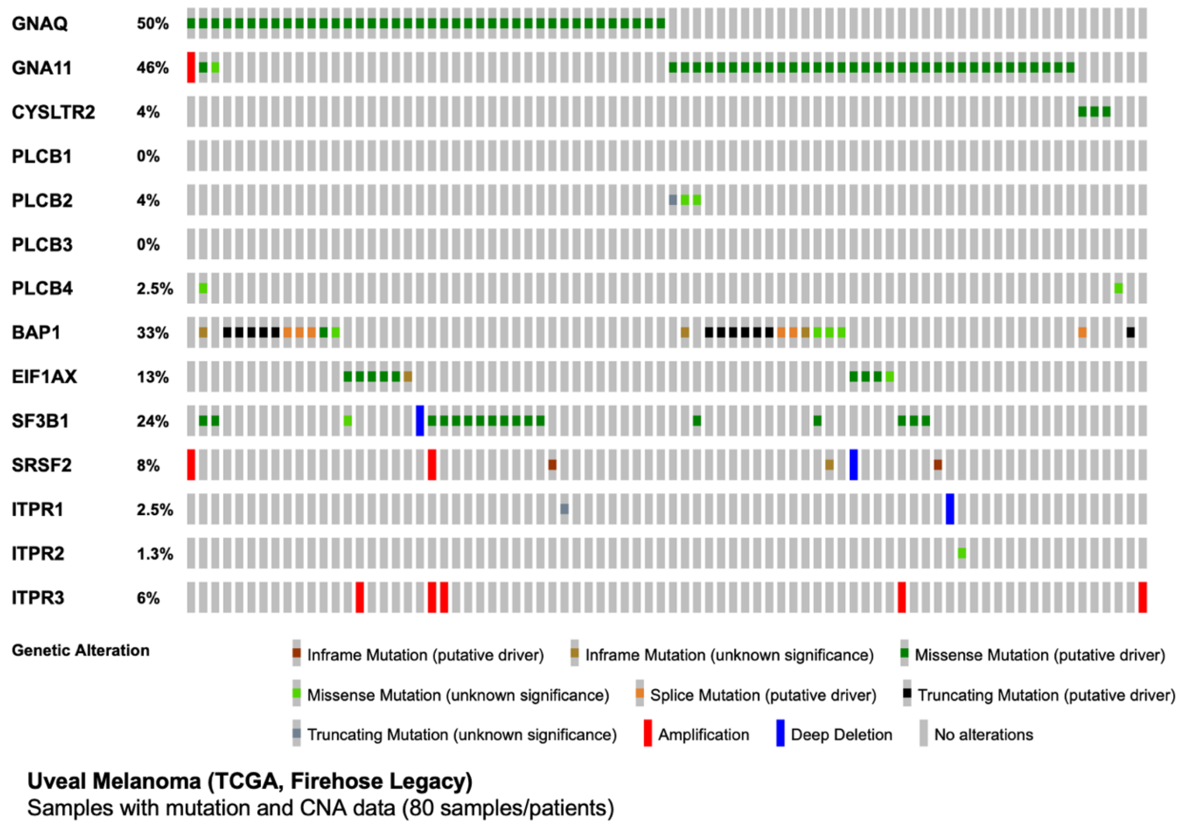

***Supplementary Figure 6. ITPR expression following G$\alpha_{q/11}$ or PKC inhibition in BRAF$^{mut}$ cells.***

**A-B** Relative *ITPR1-3* mRNA expression in the *BRAF$^{mut}$* OCM-1 cell line 24 h after treatment with the indicated concentration of either the G$\alpha_{q/11}$ inhibitor, YM-254890 (A) or the pan-PKC inhibitor, AEB071 (B) relative to the control condition. qPCR results ($2^{\Delta\Delta Ct}$ analysis) of three independent experiments are represented as mean ± SEM, with one-way ANOVA tests, not significant: ns P>0.05.

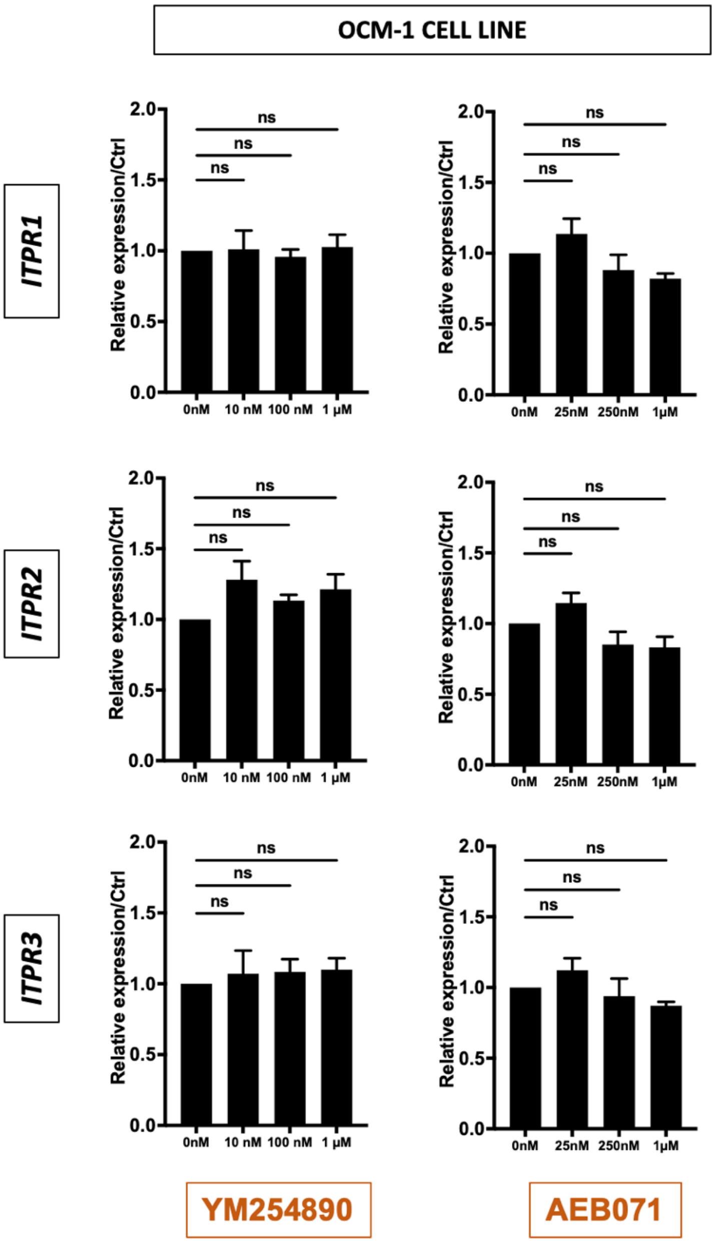

***Supplementary Movie 1. Spontaneous $Ca^{2+}$ oscillations in MP41 cell line.***

MP41 cells were probed with Fluo-4 (excitation at 494 nm and emission at 506 nm). $Ca^{2+}$ signals were recorded in the presence of 1.8 mM extracellular $Ca^{2+}$.
